# Contributions of tactile information to the sense of agency and its metacognitive representations

**DOI:** 10.1101/2023.12.15.571840

**Authors:** Angeliki Charalampaki, Anthony Buck Ciston, Elisa Filevich

**Affiliations:** Humboldt-Universität zu Berlin, Faculty of Life Sciences, Department of Psychology, Unter den Linden 6, 10099 Berlin, Germany; Bernstein Center for Computational Neuroscience Berlin, Philippstraße 13 Haus 6, 10115 Berlin, Germany; Berlin School of Mind and Brain, Humboldt-Universität zu Berlin, Luisenstraße 56, 10115 Berlin, Germany; Mind-Body-Emotion Group, Max Planck Institute for Human Cognitive and Brain Sciences, Stephanstraße 1A, D-04103 Leipzig, Germany; Hector Institute for Education Sciences & Psychology, University of Tübingen, Europastraße 6, 72072 Tübingen, Germany

**Keywords:** Sense of Agency, tactile information, motor metacognition, action-outcome, metacognition

## Abstract

We compared the contribution of tactile information to other sources of information in our representations of agency. Participants (N = 40) reached with their right hand toward a ridged plate with a specific orientation and saw online feedback that could match or differ from their action in one of three ways: the physical plate’s orientation, the action’s timing, or the hand’s position in space. Absolute subjective ratings revealed that an increased mismatch in tactile information led to a diminished sense of agency, similar to what has been reported for spatial and temporal mismatches. Further, estimations of metacognitive efficiency revealed similar M-ratios in the identification of tactile violation predictions as compared to temporal violations (but lower than spatial). These findings emphasize the importance of tactile information in shaping our experience of acting voluntarily, and show how this important component can be experimentally probed.

**Statement of Relevance:** The sense of agency is the feeling that we are the authors of our actions. It is essential not only for the control we assert over our bodies but also for how we interact with interfaces like a computer or a mobile phone. Despite the central role of touch in our daily activities, the role of tactile information in forming our sense of agency is often overlooked. In this project, we used a novel experimental design that allowed us to compare the role of tactile information relative to two other sources of information that have been previously reported to affect our agency, namely temporal and spatial information of the action. We provide evidence for the first time that tactile information is crucial for our subjective experience of agency and a tool to study this role further.

## Introduction

Being and feeling in control of one’s actions is an essential component of the self. This feeling of control that normally accompanies our voluntary movements is referred to as the sense of agency (Haggard, 2017) and has been predominantly studied in the framework of the comparator model (Frith et al., 2000). The model posits that the comparison between predicted and actual sensory consequences of an action drives potentially necessary online movement correction, and is thought to lead to the experience of agency.

One common approach to study the sense of agency experimentally is to create a mismatch in the comparison between expected and perceived visual and proprioceptive information. Participants see a stimulus external to their body (a mouse pointer on the screen (Ritterband-Rosenbaum et al., 2014; Wen et al., 2023), a virtual hand (Farrer et al., 2008; Stern et al., 2020)) that moves differently, either in time or in space, from what participants expect under conditions of complete control. Experimenters then often ask participants to rate their feelings of agency. In other instances, participants perform an action followed by an outcome (a sound (Haggard et al., 2002), a visual stimulus (Ruess et al., 2020)) and researchers measure the sense of agency using intentional binding (an implicit measure of agency) which manifests as a compression in the perceived time between a voluntary movement and its outcome (Haggard et al., 2002). These studies, together, offer compelling evidence that violations in both the temporal and spatial characteristics of visual and proprioceptive predictions decrease participants’ reported agency over their actions.

However, vision and proprioception are not the only senses involved in driving voluntary movements. Moscatelli et al. (2019) found that when participants moved their index finger over a ridged surface in the absence of visual feedback of the hand, the orientation of the ridges biased participants’ forward movements towards the orientation of the ridges. This suggests that tactile information might play a role in guiding our movements and aligns with results from the literature showing that the gating of tactile information follows an action as it unfolds, with a distortion of tactile perception before and during action execution (Colino et al., 2014) and increased sensitivity during active touch (for a review, see Juravle et al. (2017)). Thus, ignoring the potential contributions of tactile information in experimental investigations on agency neglects not only important information but also a uniquely embodied source of perceptual information.

Previous research has probed the role of tactile information in the sense of agency using the intentional binding effect. Zhao et al. (2016) found that participants did not show intentional binding following button release movements (as opposed to button presses) or following movements made without tactile feedback (moving a finger over two laser beams). Further, for both standard and touchless (under an infrared LEAP camera) button presses, adding haptic feedback (vibrotactile and mid-air touchless, respectively) led to stronger intentional binding effects relative to conditions where participants received visual feedback alone (Cornelio Martinez et al., 2017). Finally, Coyle et al. (2012) asked participants to either press a button or tap with their fingers on their contralateral arm to elicit a response. The skin-to-skin tap (recorded with a microphone) led to stronger intentional binding effects than button responses. In all cases, the results show that violations of tactile predictions can disrupt agency-related representations, emphasizing the importance of tactile information in forming the sense of agency. However, none of these studies have addressed the important question of how tactile information is incorporated — namely, whether stronger violations of tactile predictions also lead to a more salient experience of lost agency just as stronger temporal and spatial violations do. We aimed to study just that, using goal-oriented actions. Importantly, we went beyond previous research to ask not only whether the mere presence of any violation of tactile predictions leads to a loss of agency, but whether it does so in a similar fashion as other perceptual information sources.

We were interested in explicit agency ratings; but comparing the absolute value of subjective ratings across conditions might be problematic, as differences between conditions can be driven or masked by trivial response biases. Hence, to compare the subjective experience of agency across conditions, we built on a recent suggestion in the literature on agency where a standard agency task is adapted into a metacognitive task (Wang et al., 2020). Participants performed actions and discriminated which of two stimuli presented on a screen they felt more control over. Then, they reported their confidence in the correctness of their response. By doing so, it is possible to measure participants’ metacognition of agency using an estimate of metacognitive efficiency (Mratio; Maniscalco & Lau, 2012). Mratio depends on the relationship between participants’ confidence and their discrimination accuracy. Simply put, a participant with high Mratio reports high confidence when accurately assigning agency to themselves in those trials when they were in complete control — and low confidence in those where they made an agency misattribution. Conversely, a participant with low Mratio may rate with high confidence those trials where they made agency misattribution while they effectively had little control. Mratio is a measure of relative efficiency between informational levels, as it quantifies which proportion of the information available for the discrimination response is also available for the confidence judgment. It is therefore unit-free, enabling comparisons of Mratio between conditions. Unlike direct comparisons of subjective ratings, it is valid to directly compare violations of different aspects of action (spatial, temporal, and tactile) in their effect on participants’ metacognitive representations of agency.

In this study, each participant completed a standard agency and a metacognitive agency task. In each task, we included three experimental conditions: temporal, spatial, and tactile. We had two hypotheses. First, we expected that tactile prediction violations would lead to diminished subjective agency ratings. Second, we reasoned that the precision of metacognitive representations of agency following violations of different kinds could serve as a proxy for their relative importance. Hence, we expected the precision of the metacognitive representations following tactile violations to be at least as high as those following spatial and temporal violations.

### Open Practices

This study was pre-registered (osf.io/ur73j), and any deviations from the pre-registered plan are explicitly reported. The raw data, as well as R and MATLAB scripts for the analyses, are freely available at (osf.io/ur73j).

## Methods

We explored the contribution of tactile information to judgments of agency and motor metacognitive representations. Participants completed two tasks (Agency and Meta-Agency) across two sessions within a week (mean days between sessions = 2.3, SD = 2.1) where they saw their hand moving to reach their goal, in some cases with an additional spatial, temporal, or tactile manipulation. In the Agency task, we measured subjective agency ratings. In the Meta-Agency task, we measured participants’ metacognition of agency.

### Participants

We followed our pre-registered plan to collect data from a sample of n = 40 participants. We initially collected complete datasets from 47 right-handed participants (mean age = 25.6 years, SD = 4.8, 37 female, 10 male, mean handedness score = 88.6, SD = 22.1 (Oldfield, 1971)), and excluded the data from seven participants from further analyses because they met our pre-registered exclusion criterion of having very low accuracy (< 60%) in their discrimination performance on the Meta-Agency task in at least two conditions. We excluded six additional datasets from the tactile and two from the spatial condition before running the pairwise analyses because participants’ discrimination accuracy was below 60% only in these conditions. For these participants, we included only the data from the other conditions in our analyses.

Participants reported no neurological or psychiatric history, had normal or corrected-to-normal vision, and no motor impairment in either hand. They received 8 €/hour or course credits for their time. Testing took place in English. All participants signed written informed consent. The experiment was approved by the ethics committee of the Institute of Psychology of Humboldt-Universität zu Berlin (2022-57) and conducted according to the Declaration of Helsinki.

### Apparatus and tactile stimuli

For the tactile stimuli, we used two circular, ridged, 3D-resin-printed plates (60 mm Ø; Fig. 1a) centered in 70-mm-Ø holes in the top of a custom-made, 10-mm-thick wooden box. The plates were attached to the drive shafts of two stepper motors (Dual Shaft, D-cut Shaft, Nema 17, 17HM19-1684D, Step size 0.9°), which allowed the plates to rotate clockwise or counterclockwise before the beginning of each trial. The plates were flush with the surface of the box, leaving a 5 mm gap between the resin plates and the wood. The motors were controlled by two digital stepper drivers (DM332T, for Nema 17), two rotary encoders (AMT102-V, CUI Devices, Lake Oswego, OR, USA), and a microcontroller (Arduino UNO WiFi Rev2, Arduino, Monza, Italy; a description of all components and technical aspects of how to build this custom-made box can be found under: osf.io/ur73j).

**Fig. 1.**
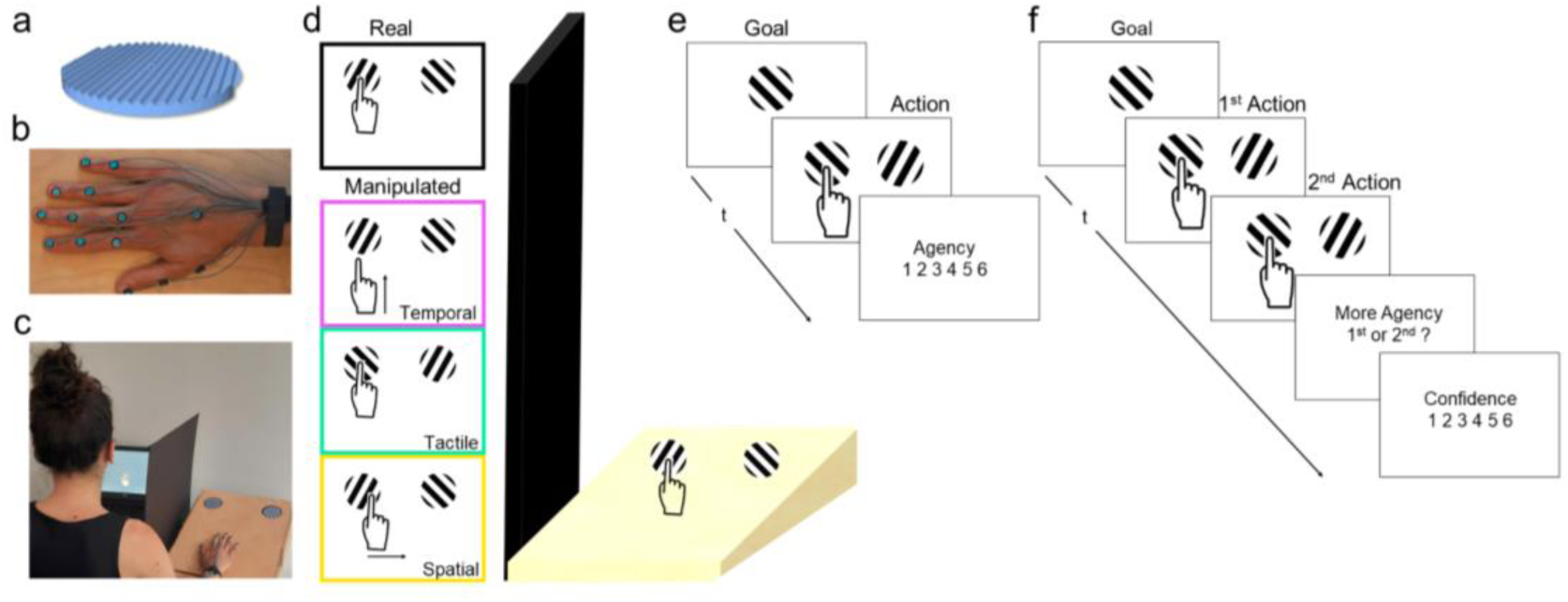
Experimental design. The tactile stimulus was a 3D-resin-printed plate (a) that rotated clockwise and counterclockwise along the vertical axis passing through its center. Participants controlled a virtual hand via fourteen sensors attached to their right hand (b). Participants faced a computer and placed their right hand behind an occluder (c). The basic motor task involved participants moving their hand and touching a specific target. The online visual feedback either matched the action participants performed or was presented with an added manipulation. For the latter, either the timing of the movement (temporal), the hand position in space (spatial), or the orientation of the virtual plates (tactile) did not match what the participants were doing or touching (d). At the beginning of each trial of the Agency task, participants saw the goal plate they had to interact with on the screen. Then, they moved their hand forward toward the plate with the same orientation as the goal and touched the center of the plate with their index finger. Participants then rated how much they felt in control of the virtual hand (e). On each trial of the Meta-Agency task, participants repeated the same action twice (moving towards and touching the target plate). The online feedback of the action was congruent with what participants did for only one of the two actions. At the end of the second action, participants reported whether they felt more in control of the virtual hand during the second action compared to the first and then rated their confidence in that decision (f).

We placed 14 sensors connected to a VIPER (Polhemus, Colchester, VT, USA) electromagnetic motion tracking system (Micro Sensor 1.8 Extra Flex™, TX2 source, and VIPER 16 SEU) on the dorsal side of participants’ right hand (Fig. 1b). These sensors tracked and recorded participants’ hand position in space (location and orientation). During the experiment, participants could not see their right hand, which was hidden behind a black occluding board. Instead, they could see only a virtual hand on the screen of a laptop (Dell Latitude 5591, a 15.6” screen: 34.42 x 19.35 cm, refresh rate 60 Hz) placed approximately 60 cm away from them and to the left of the occluder (Fig. 1c), which they could control with the sensors. The experiment was programmed in Unity (Unity Technologies, San Francisco, CA, USA) using the Unity Experiment Framework (UXF; Brookes et al., 2020).

### Motor task

Each trial started with a white fixation cross (starting position) presented in the middle of the lower part of the screen. Participants started the trial by placing their right virtual palm over the fixation cross, which then turned black. After 0.5 s the hand disappeared from the virtual display. This was to prevent participants from visually identifying manipulated intervals from the ‘jump’ in the position of the virtual hand. After the hand disappeared from the screen, a picture of a rotated ridged plate (constituting the action goal) was shown in the center of the screen for 0.5 s. 0.2 s after the offset of the goal, the two virtual plates appeared at the top of the screen in positions that corresponded to the physical positions of the real plates. Participants were tasked with using their index finger to touch the center of the plate with the same orientation as the goal. On all trials, the ridges of the two plates were rotated 45° from each other. We instructed participants to move their hand straight towards the plate (goal) and to avoid interacting with the plate at any other point aside from the center. As soon as participants placed their index finger on the center of the plate (marked with a black circle) the plates first turned blue and later turned green if participants maintained their finger on the center of the plate for the required 0.4 s. Once the plates turned green, participants could return their hand to the starting position to continue. If participants did not complete the cued action within 5 s, the plates turned red, marking the trial for exclusion. To familiarize themselves with the task and understand how much they could control the virtual hand under non-manipulated conditions, participants ran sixteen trials of the motor task alone before starting the Meta-Agency task, and eight trials prior to the Agency task.

### Experimental conditions

The online visual feedback of each action either matched what participants were doing (based on the sensor positions) and touching (orientation of physical plates), or was displayed with one of three types of manipulation: tactile, temporal, or spatial. In the tactile manipulation, the orientation of both physical plates did not match that of the virtual plates. In the temporal manipulation, the virtual hand was displayed on the screen with an added delay. Finally, in the spatial manipulation, the virtual hand deviated a number of degrees to the left or right of the participant’s real hand as measured from the starting position (Fig. 1d). We tested all three types of manipulation on separate blocks of trials on both tasks. Before each block, we instructed participants to be aware of the corresponding manipulation. That is, we explicitly pointed their attention towards either the sensation of their index finger while touching the plates, the timing of their movement, or the position of their hand in space while trying to interact with the plate with the correct orientation successfully. The reason for this was to ensure consistency among the participants. Namely, we wanted to ensure that they all had equal chances at performing well on the task.

### Meta-Agency task

On each trial of the Meta-Agency task, participants repeated the same action described in the motor task section two consecutive times (Fig. 1e). Participants first placed their hand over the white fixation cross, saw the goal for the trial, then moved toward the target plate for the first time and returned to the starting point. The plates then disappeared, and participants repeated the same action as soon as both plates reappeared on the screen. On each trial, only one of the two actions shown on the screen matched participants’ actions, whereas the other was manipulated. After completing the second action, participants used a standard computer keyboard to report whether they felt more (“F” key) or less (“A” key) in control of the second action, relative to the first. The question is analogous to asking which of the two intervals they felt more in control. Participants then rated how confident they were in their discrimination decision, on a scale from one to six, by pressing the corresponding buttons on the keyboard. Participants could press a combination of two keys to skip the trial whenever they made a procedural error. The Meta-Agency task consisted of 3 blocks of 120 trials each, one for each type of manipulation. We counterbalanced the sequence of the blocks between participants.

### Agency task

On each trial of the Agency task, participants made only one reaching movement towards the goal (Fig. 1d). As in the Meta-Agency task, the online visual feedback of the action in the Agency task could either match the real action or be manipulated. After each action, participants used the keyboard to provide a subjective agency rating by reporting on a scale from one to six how much they felt they were in control of the action. To ensure that agency ratings did not reflect participants’ uncertainty in their decision (similar to their confidence ratings in the Meta-Agency task), we emphasized that their agency ratings are subjective and that we were only interested in the level of control they experienced regardless of their confidence. We included 120 trials in each condition which were tested on separate blocks and counterbalanced between participants. Due to technical problems, one block (40 trials) from the tactile condition from one participant and one block from the spatial condition from another participant were not collected.

### Manipulated online visual feedback

For the Meta-agency task, we used separate online staircases (2-down, 1-up) for each condition to define the magnitude of manipulation used on each trial. The staircases were meant to keep participants’ discrimination accuracy at approximately 71% (Levitt, 1971). We determined the starting value of each staircase for each participant and condition separately with the training session of 16 trials. During the training session, the starting value of the tactile staircase was a 67.5° difference in orientation between the real and virtual plates, the step size was 4.5°, and the maximum possible value was 90°. The starting value of the temporal staircase was a 200 ms delay between the real and virtual hands, and the step size of the staircase was 20 ms, with no upper limit on how large the temporal delay could be. The starting value of the spatial staircase was ± 3.25° deviation, yielding a difference in position from the real hand which increased as it moved away from the fixation-cross origin. The step size of the staircase was 0.15°, with a maximum possible value set to 4.25°, as any higher value resulted in participants touching outside the edge of the plate, making the discrimination task trivial.

For each condition of the Agency task, we used five levels of magnitude of manipulation (30 trials per level of manipulation). The levels of manipulation were the same for all participants for the temporal and tactile condition — temporal (ms): [0, 50, 100, 150, 200]; and tactile (rotation degrees) [0°, 22.5°, 45°, 67.5°, 90°]. For the spatial condition, we adjusted the values two times. The reason was that, after visually inspecting the spatial staircases from the first two participants on the Meta-Agency task, we realized that the values we initially used in the Agency task (angles: [0°, 0.1°,0.2°, 0.3°,0.4°]) were not within the range of values where participants’ staircases stabilized. Therefore, we adjusted the range for the following participants so that four had the angles: [0°, 0.25°, 0.3°, 0.35°,0.4°] and the remaining 35 had angles: [0°, 3°, 3.3°, 3.6°, 3.9°]. In all cases, the manipulation magnitudes were pre-registered.

### Procedure

We tested the first two blocks of the Meta-Agency task on the first day of the experiment and the last block on the second day, prior to the Agency task. After the first training session of the Meta-agency task, participants completed an Embodiment Questionnaire (Gonzalez-Franco & Peck, 2018; Peck & Gonzalez-Franco, 2021) to measure their sense of ownership over the virtual hand.

## Data analysis

### Data exclusion

In line with our pre-registered plans, we excluded from the Meta-Agency datasets those trials in which the discrimination reaction time was under 0.2 s or above 8 s, trials marked for exclusion by the participants (e.g. error in the discrimination report), and trials where the action time was above 5 s on either interval. We deviated from the pre-registered plan and also excluded trials in which the participants touched the wrong plate on any interval. Overall, we excluded a median of 5.97 trials (IQR = 3.43 - 8.51) from each participant. Finally, after visually inspecting the staircases, we decided to deviate from the pre-registered plan by removing the first ten trials from the tactile condition from all the participants because, for many participants, the tactile staircases stabilized after the first few trials. Trial exclusion took place prior to any further data analysis.

### Confirmatory analyses

We fit linear mixed effect models using the *lme4* package (Bates et al., 2015) in R (version 4.2.2, R Core Team, 2022).

We measured participants’ metacognition of agency by estimating their metacognitive efficiency (Mratio; Maniscalco & Lau, 2012) with the maximum likelihood estimation method in R using the *metaSDT* package (Craddock, 2021). We used the *BayesFactor* package (Morey & Rouder, 2018) to get the Bayes factors for H1 (BF_10_ values), the *effectsize* package (Ben-Shachar et al., 2020) to estimate partial eta-squared, and used the Wilcoxon signed rank test for data that were not normally distributed. We report mean values and standard deviations (M = mean value ± SD).

In a set of exploratory analyses, we ran parametric and non-parametric correlations using the *ggstatsplot* package (Patil, 2021) in R.

## Results

### Agency task

We first examined participants’ responses in the embodiment questionnaire and found that participants already experienced both agency and ownership over the virtual hand (ownership score: M = 1.38 ± 0.58; agency score: M = 1.98 ± 2.24; possible range [-3,3]) after the first training session of the motor task. We then examined whether violations of tactile expectations reduce judgments of agency by measuring the relationship between agency ratings and the level of manipulation on each condition. To do so, we split the data into the three different manipulation conditions, z-transformed the magnitude of manipulation per condition (to scale the values by mean-centering to the condition-specific mean, and normalizing by the condition-specific) and fit three separate linear mixed effects models. Henceforth, we use the term “manipulation level” to refer to the ordered magnitude of the manipulation within each condition. This allowed us to align all three conditions on a single axis. Conversely, using the different absolute values of spatial, temporal, or tactile manipulation would not have allowed this.

We ran separate models per condition to avoid having a single, overly-complex model that would lead to convergence issues in the fitting procedure. We included the magnitude of manipulation as a fixed effect and by-participant random slopes and intercepts (formula: agency ∼ magnitude of manipulation + (magnitude of manipulation | participant). In line with our pre-registered hypothesis, we found a significant negative main effect of the manipulation magnitude on participants’ ratings of agency in all three conditions (Fig. 2, tactile: F(1, 39) = 99.92, p < 0.001, BF_10_ = 1.56 * 10^9^, η^2^ = 0.72, CI = [-1.01, -0.67], temporal: F(1, 39) = 273.95, p < 0.001, BF_10_ = 1.76 * 10^16^, η^2^ = 0.88, CI = [-0.87 -0.68], spatial: F(1, 39) = 76.93, p < 0.001, BF_10_ = 4.30 * 10^7^, η^2^ = 0.66, CI = [-0.49, -0.31]). These results suggest that, similar to violations of temporal and spatial predictions, an incremental mismatch between what participants touched and what they saw on the screen led to an incremental decrease in their agency ratings. This result also confirmed that the novel tactile stimuli used in our study had the expected influence on participants’ agency. Therefore, we could turn to the Meta-Agency task to test the role of tactile information on participants’ metacognitive representations of their agency judgments.

**Fig. 2.**
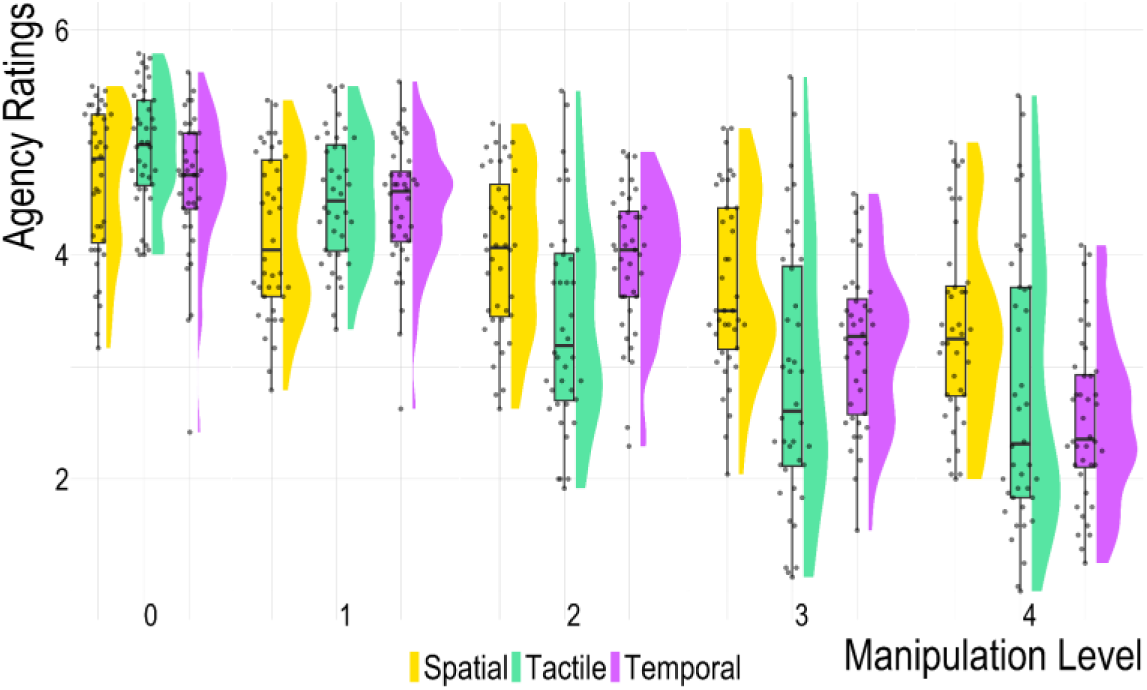
Agency Ratings relative to the level of manipulation for each of the three conditions. Each dot corresponds to one participant and condition, the boxplots represent the interquartile range and the violin plots represent the smoothed distributions of the data. In all conditions, participants rated lower agency with increasing manipulation levels. We use the term “manipulation level” to refer to the ordered magnitude of the manipulation within each condition, that could not be compared between conditions (as the manipulated dimension was different). Violations of tactile predictions resulted in lower judgments of agency, with larger manipulations leading to lower agency ratings, similar to the effect of temporal and spatial predictions.

### Meta-agency task

In the Meta-Agency task, participants made relative judgments of agency by comparing two intervals in the amount of control experienced over the virtual hand. The difference in the magnitude of manipulation between the two intervals, and hence the difficulty of the discrimination task, was titrated to each participant’s ability with an online staircase. Nevertheless, as is common in any analysis of metacognitive performance, we first verified that discrimination performance — measured with d’ — was comparable across conditions (Fig. 3). Because *d’_temporal_* values were not normally distributed, we ran the same analyses using Wilcoxon signed rank tests which confirmed the results from the pairwise t-tests (see Supplementary Material). Despite the condition-specific online staircases, participants were significantly more accurate in selecting the interval in which they were more in control in the tactile compared to the spatial condition (*d’_tactile_* = 0.96 ± 0.12; *d’_spatial_*= 0.81 ± 0.17; t(31) = -4.12, p < 0.001, Cohen’s d = -0.73, BF_10_ = 107.92). Similarly, they were significantly more accurate in the temporal compared to the spatial condition (*d’_temporal_* = 0.99 ± 0.16; *d’_spatial_* = 0.81 ± 0.17; t(37) = -5.01, p < 0.001, Cohen’s d = -0.81, BF_10_ = 1545.50). We found anecdotal evidence that participants’ discrimination performance was comparable between the tactile and temporal conditions (*d’_tactile_ =* 0.95 ± 0.14; d’_temporal_ = 1.00 ± 0.16; t(33) = 1.35, p = 0.186, Cohen’s d = - 0.23, BF_10_ = 0.42). In short, participants had worse discrimination performance in the spatial condition than in the other two conditions.

**Fig. 3.**
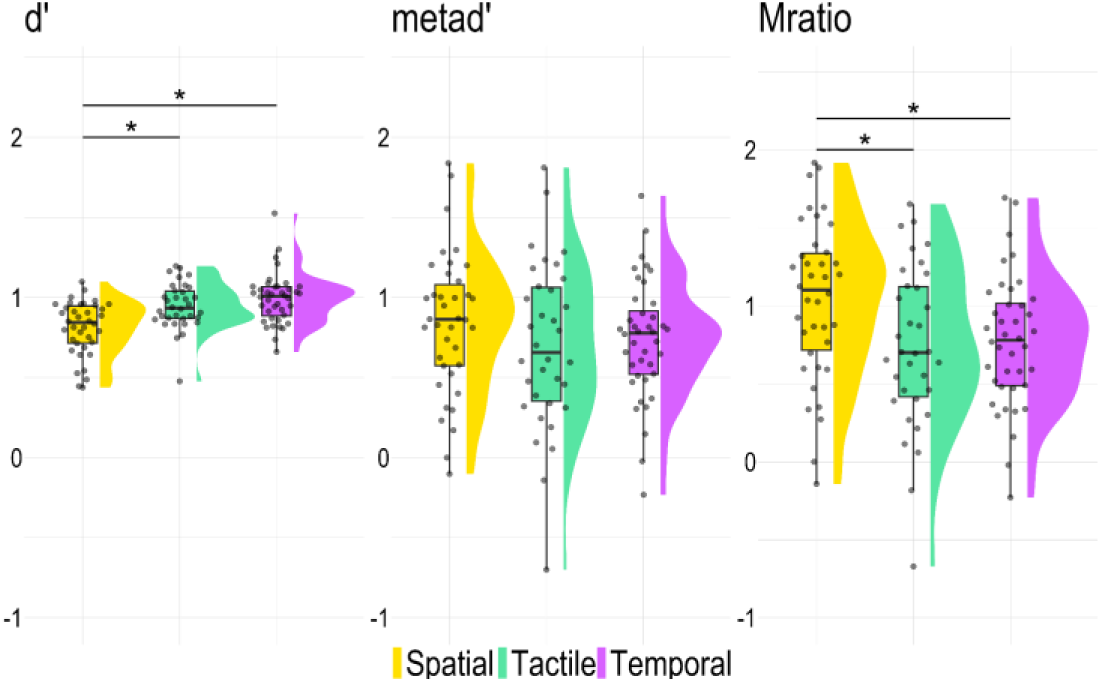
Meta-agency task. d’: Discrimination performance in the Meta-agency task. metad’: Metacognitive sensitivity. Mratio: Metacognitive efficiency. Each dot corresponds to one participant and condition, the boxplots represent the interquartile range. The violin plots represent the smoothed distributions of the data per condition. For the pairwise comparisons, we included only participants with discrimination accuracy over 60 % in the two conditions compared. The asterisks indicate that summary measures are significantly different between two conditions (*: < 0.05)

Despite the differences in discrimination performance, none of the three pairwise comparisons yielded significant differences in metacognitive sensitivity measured with meta-d’ (Fig. 3.): tactile vs. temporal (*metad’_tactile_* = 0.70 ± 0.52; *metad’_temporal_* = 0.70 ± 0.37; t(33) = 0.03, p = 0.978, Cohen’s d = 0.005, BF_10_ = 0.18), tactile vs. spatial (*metad’_tactile_* = 0.73 ± 0.52; *metad’_spatial_* = 0.83 ± 0.42; t(31) = 0.94, p = 0.353, Cohen’s d = 0.17, BF_10_ = 0.28), spatial vs. temporal (*metad’_spatial_* = 0.84 ± 0.44, *metad’_temporal_* = 0.74 ± 0.38; t(37) = 1.25, p = 0.220, Cohen’s d = 0.20, BF_10_ = 0.36).

Lastly, and crucially, we compared participants’ Mratio across conditions (Fig. 3.) to test whether participants have better metacognitive efficiency following tactile expectation violations relative to spatial and temporal ones. The results suggest this is not the case: A comparison of the tactile condition against the other two revealed that *Mratio_tactile_* was indistinguishable from *Mratio_temporal_* (*Mratio_tactile_* = 0.73 ± 0.52; *Mratio_temporal_* = 0.73 ± 0.41; t(33) = 0.06, p = 0.950, Cohen’s d = 0.01, with strong evidence against any differences BF_10_ = 0.18). Also, *Mratio_tactile_* was consistently lower than *Mratio_spatial_* (*Mratio_tactile_ =* 0.75 ± 0.53; *Mratio_spatial_* = 1.03 ± 0.51; (t(31) = 2.45, p = 0.020, Cohen’s d = 0.43, BF_10_ = 2.45). Finally, *Mratio_spatial_* was higher than *Mratio_temporal_* (*Mratio_temporal_* = 0.77 ± 0.43; *Mratio_spatial_* = 1.03 ± 0.51; t(37) = 2.76, p = 0.009, Cohen’s d = 0.45, BF_10_ = 4.53). Five participants had negative Mratio in some conditions, which indicates that they were often more confident after wrong discrimination decisions than following correct ones. To ensure that these participants’ behavior did not drive the effects we saw, we ran the same pairwise comparisons of the behavioral measures between conditions, excluding these participants from the respective comparison. The results held for the comparisons between conditions in d’, metad’, and Mratio (See Supplementary Material). Taken together, these results argue against any advantage of tactile over temporal information for participants’ metacognition of agency. Instead, the results suggest that spatial manipulations may be especially metacognitively salient relative to both tactile and temporal manipulations. Nevertheless, given the differences we found in discrimination performance, we should interpret these differences in the Mratio with caution, as Mratio has been shown to depend on d’, with higher d’ often leading to smaller Mratio values, even when the true metacognitive noise is the same (Guggenmos, 2021) despite the theoretical expectations that it will not (Maniscalco & Lau, 2012).

### Correlations between measures of agency

In line with our pre-registered plans, we investigated the relationship between the subjective agency ratings and metacognitive judgments of agency. To do so, we first extracted the random slope corresponding to each participant from the condition-specific linear mixed-effects models (see Agency task above). These slopes for each condition represent how sensitive participants’ judgments of agency are to the incremental mismatches between their perceptual predictions and observations. Then, for each condition, we ran non-parametric correlations (because the individual agency slopes were not normally distributed) between the agency slopes and the corresponding Mratio. For all conditions, we found a negative relationship between the two measures (Fig. 4.), with participants with larger Mratio values (indicating higher metacognitive efficiency) also generally showing steeper agency slopes (indicating higher sensitivity in the magnitude of manipulation). Nevertheless, none of the correlations between these two summary measures was significant in any of the conditions (tactile: S = 8640, Spearman’s rho-hat = -0.32, p = 0.060, CI = [-0.60, 0.03], n = 34; spatial: S = 11216, Spearman’s rho-hat = -0.23, p = 0.170, CI = [-0.52, 0.11], n = 38; temporal: S = 11866, Spearman’s rho-hat = -0.11, p = 0.490, CI = [-0.42, 0.21], n = 40). These results suggest that while related, Agency slope and Mratio do not result from the same underlying processes.

**Fig. 4.**
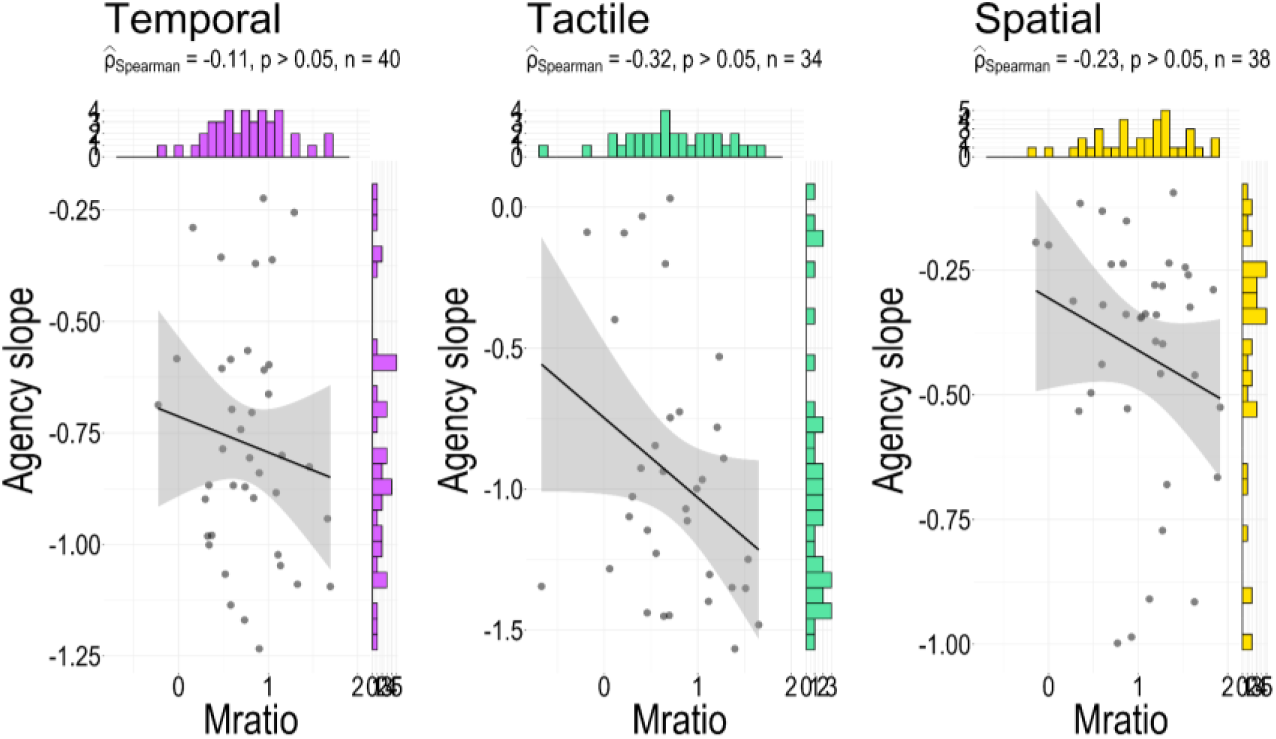
Correlation analyses between measures of agency: Agency slope and Mratio. We found no significant correlations between these two measures in any of the conditions. In all panels, each dot corresponds to one participant, the histograms indicate the distribution of data for each measure, the black lines represent the regression line and the shaded area corresponds to the 95% confidence interval.

## Exploratory analyses

### Correlations between sensitivity across conditions

In a set of exploratory analyses, we first tested how the agency measures relate between the different conditions. We found a significant positive correlation only between the slopes of the spatial and temporal condition (spatial vs. temporal: S = 7214, Spearman’s rho-hat = 0.32, p = 0.042, CI = [0.004, 0.58], n = 40; tactile vs. temporal: S = 7770, Spearman’s rho-hat = 0.27, p = 0.091, CI = [-0.05, 0.54], n = 40; spatial vs. tactile: S = 7728, Spearman’s rho-hat = 0.28, p = 0.086, CI = [-0.05, 0.55], n = 40; Fig. 5a.). Then, we tested for possible correlations between the Mratio values obtained from each condition. We found a significant correlation between the Mratios of only the tactile and the temporal conditions (tactile-temporal: t(34) = 3.27, Pearsons’s r = 0.50, p = 0.003, CI = [0.19,0.72], BF_10_ = 18.14, n = 34; tactile-spatial: t(32) = 1.44, Pearsons’s r =0.25, p = 0.160, CI = [-0.10, 0.55], BF_10_ = 0.67, n = 32; spatial-temporal: t(38) = 1.39, Pearsons’s r = 0.22, p = 0.170, CI = [-0.10, 0.51], BF_10_ = 0.59, n = 38; Fig. 5b). This correlation between *Mratio_tactile_* and *Mratio_temporal_* suggests that temporal and tactile information contribute through similar mechanisms to participants’ metacognitive representations of agency.

**Fig. 5.**
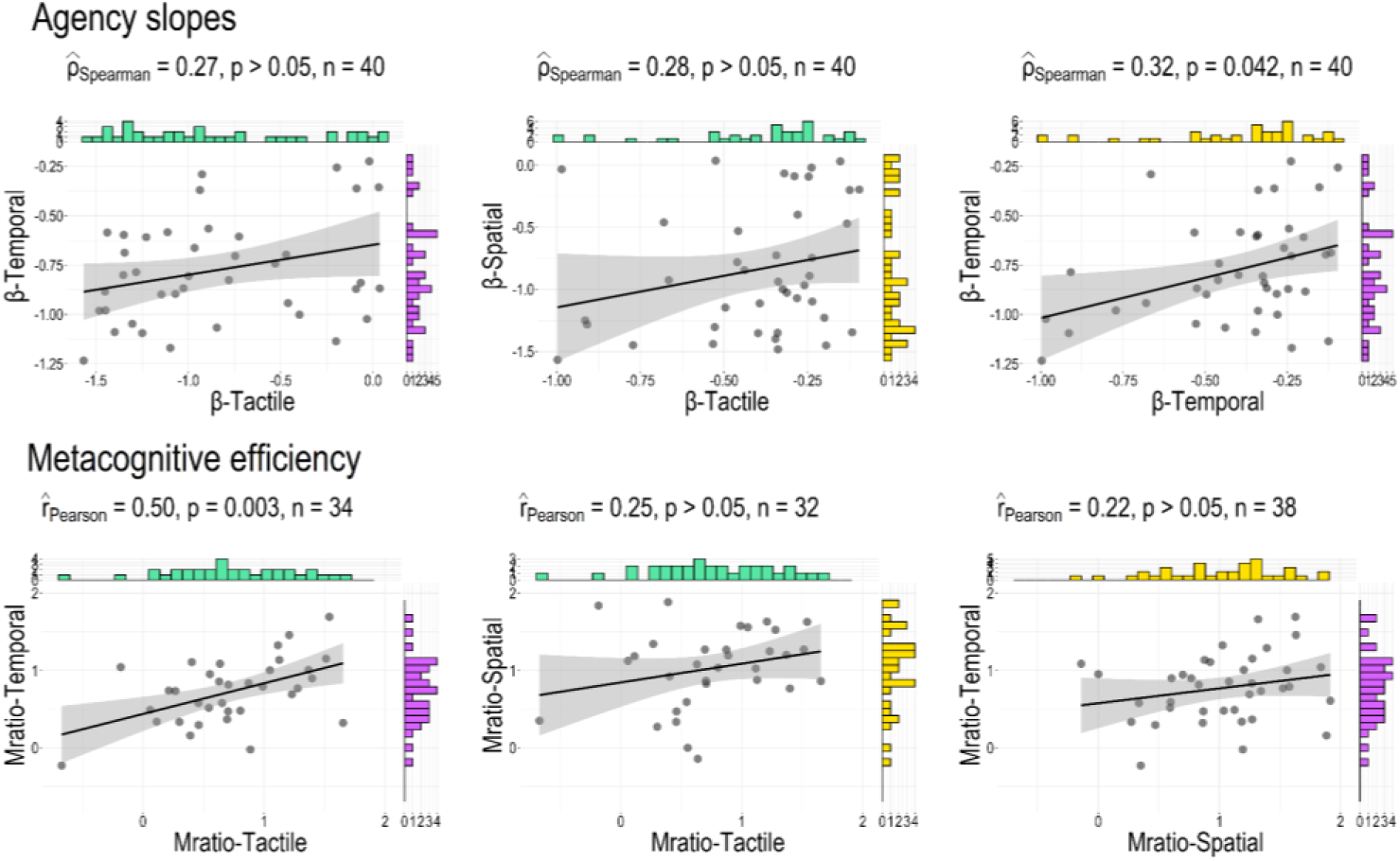
Correlation analyses of different measures of agency sensitivity between the pairs of conditions. Significant correlations were found between *agency slope_temporal_* and *agency slope_spatial_* and between *Mratio_tactile_* and *Mratio_temporal_*. In all panels, each dot corresponds to one participant, the histograms indicate the distribution of data for each condition, the black lines represent the regression line and the shaded area corresponds to the 95% confidence interval.

### Discrimination decision and confidence biases

A 2IFC design, like the one we used, has been suggested to lead to smaller response biases than the equivalent 2AFC discrimination tasks (Green & Swets, 1974; Kingdom & Prins, 2016; Macmillan & Creelman, 2004), but response biases have still been reported (Yeshurun et al., 2008). We therefore tested for response and confidence biases in all three conditions. First, we obtained participants’ decision criteria from the output of the *metaSDT* package, which implements the standard formula to estimate the criterion as: c = -0.5 * [z(hit rate) + z(false alarm)], where z is the normal cumulative distribution function. Participants’ decision criteria were significantly different from zero in all three conditions (tactile: t(33) = 4.82, p < 0.001, Cohen’s d = 0.82, BF_10_ =729.50; spatial: t(37) = 6.10, p < 0.001, Cohen’s d = 0.98, BF_10_ = 3.13*10^4^; temporal: t(39) = 7.60, p < 0.001, Cohen’s d = 1.20, BF_10_ = 3.79 *10^6^). Thus, despite the 2IFC task, participants selected the second interval as the one they were more in control more often than was presented (in 56 % of the tactile trials, 58 % of the spatial trials and 60 % of the temporal trials when, in reality, they were in control of the action in the second interval in all three conditions in 49.9 % of the trials).

We then tested for potential biases in participants’ confidence. We fit a linear mixed-effect model that included condition, manipulated interval, interval selected and scaled magnitude of manipulation as main effects, along with the interaction between the condition, manipulated interval, the interval-selected, and with by-participant random intercept (formula: confidence ∼ condition * interval-manipulated * interval-selected + manipulation magnitude + (1 | participant)). Frequentist statistics revealed a significant three-way interaction between the condition, manipulated interval, and interval selected (F(1,12672) = 3.23; p = 0.039; Fig. 6), albeit with a small effect size (η^2^ = 0.0005). While Bayesian hypothesis testing revealed evidence against an effect of the three-way interaction term (BF_10_ = 0.002). Frequentist and Bayesian statistics point towards different conclusions, but we honored our pre-registered plans to interpret both. We therefore ran post-hoc tests, using the *emmeans* package (Lenth 2023) in R, to tease it apart (reported in detail in the Supplementary Material). These revealed that participants were more confident when they were correct. Additionally, participants tended to report higher confidence when they selected the second interval, in all conditions when this response was incorrect, and only in the tactile and temporal conditions, for both correct and incorrect responses. We also found a significant main effect of manipulation magnitude (F(1, 12671) = 140.30; p < 0.001; BF_10_ = 1.88*10^28^; η^2^ = 0.01). The latter indicates that a larger level of manipulation resulted in higher confidence ratings. Given that we also found a response bias (participants selected the second interval more often), we also measured participants’ metacognitive efficiency by fitting the data using response-specific meta-d models (Maniscalco & Lau, 2014). We found no significant differences in participants’ Mratio for selecting the first or the second interval in any conditions (See Supplementary Material). Therefore, the results from the confirmatory analysis hold despite participants’ biases.

**Fig. 6.**
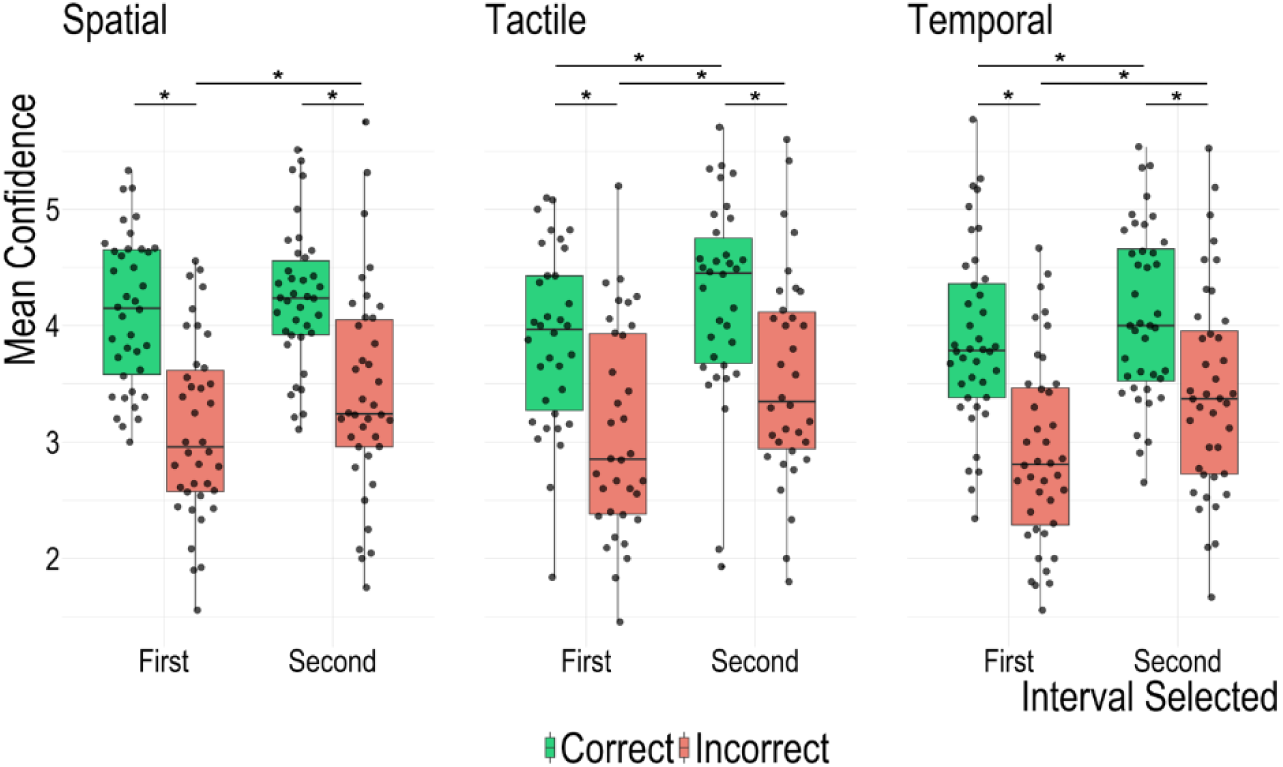
Mean confidence ratings split according to the interval participants selected as the one in which they had more control and the accuracy of that decision for all three conditions. Each dot corresponds to one participant, and the boxplots represent the interquartile range. We found a three way-interaction effect on confidence between the condition, manipulated interval, and interval selected. We present the most relevant results from post-hoc pairwise comparisons between the factors while adjusting for multiple comparisons using the Bonferroni correction. The figure illustrates the effects reported in the text: Participants were more confident when they were correct. In the tactile and temporal conditions, participants were more confident when they selected the second interval regardless of their accuracy. In the spatial condition this was only true when participants wrongly selected the second interval. (*: < 0.05)

## Discussion

This paper delved into the role of tactile information in explicit agency judgments and their metacognitive representations. Participants moved their hand toward a target textured surface, and we manipulated the visual representation of their moving hand, allowing us to study their embodied sense of agency (Christensen & Grünbaum, 2017; Wen, 2019). In two separate tasks (Agency and Meta-Agency) we measured subjective judgments and metacognitive representations of agency and studied the relationships between them. Briefly put, participants’ responses in the Agency task revealed that violations of tactile predictions reduce the reported sense of agency, much like violations of temporal and spatial predictions (typically used in the literature). Additionally, participants’ responses in the Meta-Agency task revealed that their metacognitive access to representations of (loss of) agency was above-chance, in line with previous studies (Krugwasser et al., 2019, 2022; Penton et al., 2022; Stern et al., 2020; Wang et al., 2020; Wen et al., 2023). Finally, comparing participants’ two kinds of ratings (agency and confidence in agency discrimination) suggested that the two measures might differ, also in agreement with previous work (Constant et al., 2022). We will now elaborate on these three complementary findings.

In our Agency task, we saw that tactile information plays a role in subjective agency ratings. This finding complements recent accounts highlighting the significance of tactile information in guiding motor control (Moscatelli et al., 2019). In our task, both the movement and the outcome were tightly linked to participants’ body. Arguably, therefore, the Agency task required judgments about embodied feelings of agency (over the movement or the outcome) — more so than other studies on external agency where judgments are about distal outcomes associated with movements (Christensen & Grünbaum, 2017; Wen, 2019). Therefore, our design and the tactile stimuli allow us to bridge embodied and external agency, which have thus far been studied in separation.

We found that metacognitive efficiency (Mratio) was comparable between the temporal and tactile conditions and that metacognitive efficiency correlated positively between the tactile and temporal conditions. Together, these results suggest that tactile information contributes to the experience of agency in a comparable manner, and perhaps through similar mechanisms, as temporal information does. While relatively straightforward to control experimentally, temporal manipulations have recently been criticized on conceptual grounds. Wen (2019) argued that temporal delays might not have the expected effect on participants’ agency because, for instance, a delay in feedback may impair participants’ ability to monitor and continuously correct a movement. On the other hand, tactile manipulations do not suffer from this conceptual limitation and, given the apparent practical equivalence to temporal manipulations that we reported here, might be a promising alternative for experimental manipulations.

We also found that M-ratios for tactile and temporal violations were consistently lower than spatial ones, implying noisier metacognitive representations. Two non-exclusive explanations can account for this result. First, is it possible that this is caused by an estimation error: Despite the online adaptive staircases, discrimination performance (d’) was consistently lower in the spatial condition than in the other two conditions. Mratio is, in theory, independent of discrimination performance (Maniscalco & Lau, 2012), but in practice a dependency exists (Guggenmos, 2021; Rausch et al., 2023). Because Mratio increases slightly with decreasing d’, this could explain (partially, at least) both the consistently higher Mratio values in the spatial condition and the absence of correlations with Mratio in the other conditions. Therefore, we interpret this difference with caution. Notwithstanding this potentially more parsimonious explanation, previous studies have shown that participants tend to recognize temporally manipulated feedback to be a delayed version of their movements (Farrer et al., 2008) and more often attribute temporally manipulated feedback to themselves than spatially manipulated feedback (Stern et al., 2020) but see (Krugwasser et al., 2019)). Additionally, we note a limitation of our design: In cases where the spatial manipulation was large enough, participants could have felt the edge of the plate but seen their fingers touching only the plate. We aimed to avoid this situation by setting a maximum to the spatial staircase of exactly the radius of the plate, but this may not have been enough. Hence, in these trials, participants would have had both spatial and tactile information to guide their judgments. We expect this to have affected both agency and Meta-Agency judgments, and so it calls for caution against strongly interpreting an advantage of spatial manipulations.

### 2IFC-metacognitive paradigms to study agency

Recently studies have shifted away from simple agency ratings and have instead incorporated agency judgments in metacognitive tasks (Krugwasser et al., 2019, 2022; Penton et al., 2022; Stern et al., 2020; Wang et al., 2020; Wen et al., 2023). This aims to explicitly quantify the uncertainty in participants’ judgments, thereby providing a bias-free measure of participants’ agency representations. While attractive, this approach entails two potential limitations. First, as we saw in our study, response and confidence biases are still present — overall, we found decision and confidence biases in all three conditions as participants tended to select the second interval as the one in which they felt more in control and report higher confidence when they did. Additionally, and more conceptually, in line with a recent study showing that agency judgments do not rely on the same computations as confidence ratings do (Constant et al., 2022), we found a negative but non-significant relationship between the subjective and metacognitive representations of agency in all three conditions. Therefore, care should be taken before simply replacing agency judgments with 2IFC-confidence tasks in the study of agency.

### Conclusion

This study shows that tactile predictions, like better-studied temporal and spatial predictions, contribute to participants’ both subjective and metacognitive representations of agency. Moving forward, our experimental design can be further developed to address, for example, how different types of predictions computationally combine to create a unified experience of agency.

## Acknowledgments

We thank Lorenzo Malloni for help with experiment materials and Lucie Charles for helpful comments on a previous version of the manuscript.

## Author Contributions

Angeliki Charalampaki: Conceptualization; Methodology; Formal analysis; Investigation; Data Curation; Writing – Original Draft; Writing – Review & Editing; Visualization, Validation.

Elisa Filevich: Conceptualization; Methodology; Formal analysis; Resources; Writing – Original Draft; Writing – Review & Editing; Writing – Review & Editing; Supervision; Funding Acquisition; Validation. Anthony Ciston: Software; Methodology, Writing – Review & Editing.

