## Supplementary Material for "Contributions of tactile information to the sense of agency and its metacognitive representations"

***Supplementary material to the confirmatory analysis***

Meta-Agency Task

*Non-parametric tests of d' differences*

The Wilcoxon signed rank tests on the differences between d' in all pairs of comparisons confirmed the results from the pairwise t-tests: tactile-temporal: W= 482, $p = 0.240$, $r = 0.21$; tactile-spatial: W = 250, $p < 0.001$, $r = 0.60$; temporal-spatial: W = 302, $p < 0.001$, $r = 0.66$).

*Excluding participants with Mratio < 0*

Given that five participants had negative Mratio in at least one of the three conditions, we ran the pairwise comparisons of the behavioral measures between the conditions, excluding these participants from the respective tests. Similar to the results we report in the main text, participants' discrimination performance was significantly lower in the spatial ($d'_{spatial} = 0.82 \pm 0.16$) compared to the tactile condition ($d'_{tactile} = 0.97 \pm 0.12$; $t(28) = -4.13$, $p < 0.001$, Cohen's d = -0.77, $BF_{10} = 99$) and also lower than in the temporal condition ($d'_{spatial} = 0.82 \pm 0.17$; $d'_{temporal} = 1.00 \pm 0.16$; $t(34) = -4.64$, $p < 0.001$, Cohen's d = -0.78, $BF_{10} = 470.46$). Discrimination performance was not different between the tactile ($d'_{tactile} = 0.95 \pm 0.15$) and temporal conditions ($d'_{temporal} = 0.99 \pm 0.17$; $t(30) = -1.18$, $p = 0.247$, Cohen's d = -0.21, $BF_{10} = 0.36$). Non-parametric analysis confirmed the results (tactile-temporal: Wilcoxon Signed-Ranks test: W= 414, $p = 0.355$, $r = 0.18$; tactile-spatial: W = 199, $p < 0.001$, $r = 0.63$; temporal-spatial: W = 267, $p < 0.001$, $r = 0.65$).

In line with the comparisons of participants' metacognitive sensitivity (metad') that we report in the main text, we found that after excluding these five participants, metad' was not different between any of the conditions ($metad'_{tactile} = 0.77 \pm 0.45$ versus $metad'_{temporal} = 0.74 \pm 0.31$: $t(30) = 0.34$, $p = 0.739$, Cohen's d = 0.06, $BF_{10} = 0.20$; $metad'_{tactile} = 0.81 \pm 0.44$ versus $metad'_{spatial} = 0.85 \pm 0.35$: $t(28) = 0.41$, $p = 0.688$, Cohen's d = 0.08, $BF_{10} = 0.21$; $metad'_{spatial} = 0.88 \pm 0.41$ versus $metad'_{temporal} = 0.79 \pm 0.34$: $t(34) = 1.22$, $p = 0.232$, Cohen's d = 0.21, $BF_{10} = 0.36$).

Finally, in agreement with the results we report in the main text, excluding these five participants did not change the conclusions from the pairwise comparisons between Mratio between conditions. Namely, we found strong evidence for higher Mratio only in the spatial condition relative to the temporal condition ($Mratio_{spatial}$ = 1.09 ± 0.47 vs. $Mratio_{temporal}$ = 0.82 ± 0.39; t(34) = 3.01, p = 0.005, Cohen's d = 0.51, $BF_{10}$ = 7.84) and relative to the tactile condition ($Mratio_{spatial}$ = 1.06 ± 0.44 vs. $Mratio_{tactile}$ = 0.84 ± 0.45; t(28) = 2.31, p = 0.028, Cohen's d = 0.43, $BF_{10}$ = 1.92). However, we found strong evidence against any difference in the Mratio of the tactile compared to the temporal condition ($Mratio_{tactile}$ = 0.80 ± 0.45; $Mratio_{temporal}$ = 0.77 ± 0.37: t(30) = 0.41, p = 0.683, Cohen's d = 0.07, $BF_{10}$ = 0.21). Together, these results confirm those we report in the main text.

*Response-specific Metad'*

To test how the response and confidence biases affect participants' metacognitive efficiency (Mratio), we ran a response-specific metad' analysis (Maniscalco & Lau, 2014) in R and Matlab (R2020a, MathWorks, Natick, MA) and. We first split the trials for each separate condition according to whether participants selected the first or second interval. We will call these $Response_{First}$ and $Response_{Second}$. We then compared the Mratio between $Response_{First}$ and $Response_{Second}$ for each condition. Given that $Mratio_{Second}$ in the spatial condition was not normally distributed, we compared $Mratio_{First}$ and $Mratio_{Second}$ in this condition using Wilcoxon signed rank tests. We found that there was no significant difference between the Mratio for the $Response_{First}$ and $Response_{Second}$ trials for any of the conditions, and in some cases there was evidence for the null hypothesis of no difference (tactile: $Mratio_{First}$ = 0.72 ± 0.74, $Mratio_{Second}$ = 0.74 ± 0.47, t(33) = 0.25, p = 0.801, Cohen's d = 0.04, $BF_{10}$ = 0.19; spatial: $Mratio_{First}$ = 1.33 ± 0.81, $Mratio_{Second}$ = 0.95 ± 0.62, W = 543, p = 0.063, r = 0.37; temporal: $Mratio_{First}$ = 0.87 ± 0.69, $Mratio_{Second}$ = 0.71 ± 0.42, t(39) = -1.33, p = 0.192, Cohen's d = -0.21, $BF_{10}$ = 0.38).

Further, we compared $Response_{First}$ and $Response_{Second}$ between conditions. We ran non-parametric analysis for any pair of comparisons that included $Mratio_{Second\text{-}spatial}$, which was not normally distributed. For $Response_{First}$, we found that Mratios were significantly higher in the spatial condition compared to both the tactile and temporal conditions ($Mratio_{First\text{-}tactile}$ = 0.76 ± 0.73 vs. $Mratio_{First\text{-}spatial}$ = 1.39 ± 0.79: t(31) = 3.58,

$p = 0.001$, Cohen's $d = 0.63$, $BF_{10} = 28.60$; $Mratio_{First\text{-}temporal} = 0.86 \pm 0.69$ vs. $Mratio_{First\text{-}spatial} = 1.33 \pm 0.81$: $t(37) = 2.88$, $p = 0.007$, Cohen's $d = 0.47$, $BF_{10} = 5.90$).

The same analysis of Mratios for $Response_{Second}$ revealed that the spatial condition had higher response-specific Mratios than that of the temporal condition ($Mratio_{Second\text{-}temporal} = 0.73 \pm 0.43$ vs. $Mratio_{Second\text{-}spatial} = 0.95 \pm 0.62$: $W = 931$, $p = 0.030$, $r = 0.48$). The rest of the comparisons resulted in no significant differences between Mratios ($Mratio_{First\text{-}tactile} = 0.72 \pm 0.74$ vs. $Mratio_{First\text{-}temporal} = 0.81 \pm 0.63$: $t(33) = -0.75$, $p = 0.460$, Cohen's $d = -0.13$, $BF_{10} = 0.24$; $Mratio_{Second\text{-}tactile} = 0.74 \pm 0.47$ vs. $Mratio_{Second\text{-}temporal} = 0.68 \pm 0.42$: $t(33) = 0.71$, $p = 0.483$, Cohen's $d = 0.12$, $BF_{10} = 0.23$; $Mratio_{Second\text{-}tactile} = 0.74 \pm 0.46$ vs. $Mratio_{Second\text{-}spatial} = 0.91 \pm 0.63$: $W = 624$, $p = 0.135$, $r = 0.26$).

We see that while response-specific metacognitive efficiency was overall lower in both intervals in the temporal condition relative to the spatial condition, response-specific metacognitive efficiency in the tactile condition was lower for only the first interval. These results broadly confirm those reported in the main text.

*Post-hoc analyses of the linear mixed-effect models of confidence*

Post-hoc analyses revealed that in all conditions participants were more confident when they were correct compared to when they were wrong (tactile: Mean ($Confidence_{Correct\ Agency\ First} - Confidence_{Wrong\ Agency\ First}$) = 0.26, SE = 0.07, $p < 0.001$; Mean ($Confidence_{Correct\ Agency\ Second} - Confidence_{Wrong\ Agency\ Second}$) = 1.10, SE =0.08, $p < 0.001$; temporal: Mean ($Confidence_{Correct\ Agency\ First} - Confidence_{Wrong\ Agency\ First}$) = 0.39, SE = 0.06, $p < 0.001$; Mean ($Confidence_{Correct\ Agency\ Second} - Confidence_{Wrong\ Agency\ Second}$) = 1.10, SE = 0.07, $p < 0.001$; spatial: $Confidence_{Correct\ Agency\ First} - Confidence_{Wrong\ Agency\ First}$ = 0.61, SE =0.06, $p < 0.001$; Mean ($Confidence_{Correct\ Agency\ Second} - Confidence_{Wrong\ Agency\ Second}$) = 1.10, SE = 0.07, $p < 0.001$). Post-hoc analyses revealed also that in both tactile and temporal conditions, but not in the spatial condition, participants were more confident when they correctly selected the second interval as the one where they were more in control (tactile: $p < 0.001$, Mean ($Confidence_{Correct\ Agency\ Second} - Confidence_{Correct\ Agency\ First}$) = 0.35, SE = 0.05; temporal: $p = 0.003$, Mean ($Confidence_{Correct\ Agency\ Second} - Confidence_{Correct\ Agency\ First}$) = 0.20, SE = 0.05;

spatial: $p = 1.000$, Mean ($\text{Confidence}_{\text{Correct Agency Second}}$ - $\text{Confidence}_{\text{Correct Agency First}}$) = -0.03, SE = 0.05). Participants were also more confident when they were wrong in selecting the second interval as the one where they were more in control in all three conditions compared to when they were wrong and they selected the first (tactile: $p < 0.001$, Mean ($\text{Confidence}_{\text{Wrong Agency Second}}$ - $\text{Confidence}_{\text{Wrong Agency First)}}$ = 0.48, SE = 0.09; temporal: $p < 0.001$, Mean ($\text{Confidence}_{\text{Correct Agency Second}}$ - $\text{Confidence}_{\text{Wrong Agency First}}$) = 0.53, SE = 0.08; spatial:$p < 0.001$, Mean ($\text{Confidence}_{\text{Correct Agency Second}}$ - $\text{Confidence}_{\text{Wrong Agency First}}$) = 0.37, SE = 0.08). Further, participants' confidence was higher in the spatial compared to the temporal condition when participants were correct in general (Mean ($\text{Confidence}_{\text{Correct Agency First spatial}}$ - $\text{Confidence}_{\text{Correct Agency First temporal}}$) = 0.27, SE = 0.05, $p < 0.001$; Mean ($\text{Confidence}_{\text{Correct Agency Second spatial}}$ - $\text{Confidence}_{\text{Correct Agency Second temporal}}$) = 0.17, SE = 0.04, $p = 0.016$). Confidence was also higher in the spatial compared to the tactile condition but only when participants correctly selected the first interval (Mean ($\text{Confidence}_{\text{Correct Agency First spatial}}$ - $\text{Confidence}_{\text{Correct Agency First tactile}}$) = 0.23, SE = 0.05, $p < 0.001$; Mean ($\text{Confidence}_{\text{Correct Agency Second spatial}}$ - $\text{Confidence}_{\text{Correct Agency Second tactile}}$) = -0.03, SE = 0.05, $p = 1.000$). On the contrary, confidence was higher in tactile relative to temporal trials only when participants were correct in selecting the second interval (Mean ($\text{Confidence}_{\text{Correct Agency Second tactile}}$ - $\text{Confidence}_{\text{Correct Agency Second temporal}}$) = 0.20, SE = 0.05, $p = 0.002$; Mean ($\text{Confidence}_{\text{Correct Agency First tactile}}$ - $\text{Confidence}_{\text{Correct Agency First temporal}}$) = 0.05, SE = 0.05, $p = 1.000$). Finally, participants' confidence ratings did not differ between conditions when participants were wrong, regardless of the interval that was manipulated.

In addition to the three-way interaction, we also found a significant interaction between condition and interval-selected ($F(1, 12674) = 4.10$; $p = 0.017$; $BF_{10} = 0.02$; $\eta^2_P = 0.0006$); a significant interaction between interval-selected and manipulated-interval ($F(1, 12672) = 725.20$; $p < 0.001$; $BF_{10} = 8.38*10^{155}$; $\eta^2_P = 0.05$); a significant main effect of condition ($F(1, 12685) = 16.70$; $p < 0.001$; $BF_{10} = 2.40*10^{6}$; $\eta^2_P = 0.003$); a significant main effect of interval-selected ($F(1, 12680) = 143.80$; $p < 0.001$; $BF_{10} = 2.86*10^{8}$; $\eta^2_P = 0.01$); a significant main effect of manipulated-interval ($F(1, 12673) = 20.20$; $p < 0.001$; $BF_{10} = 0.07$; $\eta^2_P = 0.002$); and

***Supplementary exploratory analyses***

*Mean Confidence ratings*

When we evaluated participants' confidence ratings in their discrimination decisions, we only found anecdotal evidence for a difference in the mean confidence ratings between the spatial and temporal conditions ($Confidence_{spatial}$ = 3.94 ± 0.59; $Confidence_{temporal}$ = 3.78 ± 0.72; t(37) = 2.04, p = 0.048, Cohen's d = 0.33, $BF_{10}$ = 1.11). Mean confidence ratings were higher in the spatial compared to the temporal condition, (in line with Stern et al.,2020)). Mean confidence ratings in the tactile condition were not significantly different from those in the temporal or spatial condition ($Confidence_{tactile}$ = 3.92 ± 0.77 vs. $Confidence_{temporal}$ = 3.77 ± 0.68; t(33) = 1.63, p = 0.112, Cohen's d = 0.28, $BF_{10}$ = 0.61; $Confidence_{tactile}$ = 3.89 ± 0.79 vs.; $Confidence_{spatial}$ = 3.89 ± 0.62; t(31) = 0.02, p = 0.984, Cohen's d = 0.003, $BF_{10}$ = 0.19)). We then compared the mean confidence ratings excluding the five participants with negative Mratio. We found anecdotal evidence for a difference between mean confidence ratings in the tactile condition relative to the temporal condition ($Confidence_{tactile}$ = 3.98 ± 0.68; $Confidence_{temporal}$ = 3.80 ± 0.65; t(30) = 2.06, p = 0.048, Cohen's d = 0.37, $BF_{10}$ = 1.21). However, we found no evidence for a difference between the mean confidence ratings in the tactile condition relative to the spatial or temporal conditions ($Confidence_{tactile}$ = 3.90 ± 0.69 vs. $Confidence_{spatial}$ = 3.82 ± 0.56; t(28) = -0.79, p = 0.436, Cohen's d = -0.15, $BF_{10}$ = 0.26; $Confidence_{spatial}$ = 3.88 ± 0.54 vs. $Confidence_{temporal}$ = 3.74 ± 0.70; t(34) = 1.75, p = 0.090, Cohen's d = 0.30, $BF_{10}$ = 0.71).

*d' and Mratio correlations*

We also explored the relationship between d' and Mratio within each condition. We found that there was a significant negative correlation only in the temporal condition (tactile: S = 6220, Spearman's rho-hat = 0.05, p = 0.780, CI = [-0.3,0.39], n = 34; spatial: S = 9930.01, Spearman's rho-hat = -0.09, p = 0.610, CI = [-0.4, 0.25], n = 38; temporal: S = 15048, Spearman's rho-hat = -0.41, p = 0.008, CI = [-0.65, -0.11], n = 40; Fig. S1). While not expected in theory, this dependency has been empirically shown and discussed before (Guggenmos, 2021).

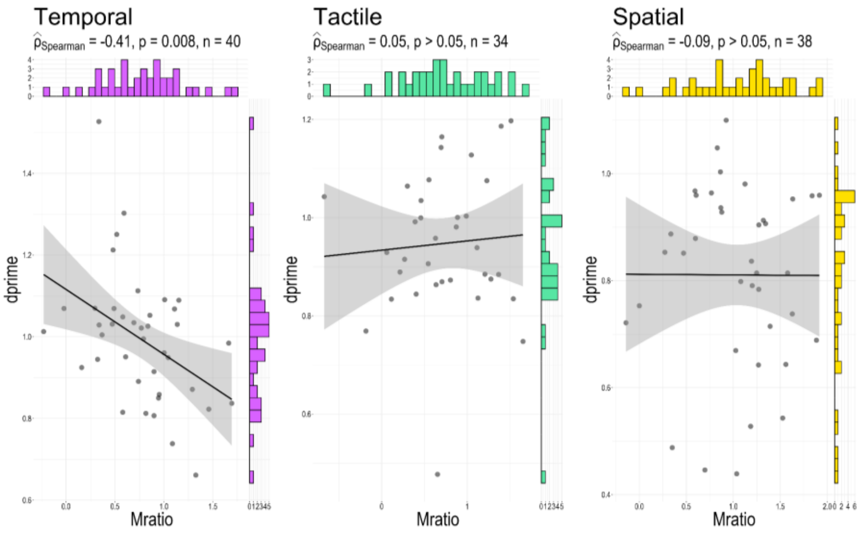


**Fig. S1.** Correlation between the d' (discrimination performance) and Mratio (metacognitive efficiency) from the Meta-Agency task. Significant correlations were found only between the d' and Mratio in the temporal condition. In all panels, each dot corresponds to one participant, the histograms indicate the distribution of data for each condition, the black lines represent the regression line and the shaded area corresponds to the 95% confidence interval.

**Embodiment Questionnaire:**

The questions used in the Embodiment Questionnaire:

1. I felt as if the virtual hand was my hand
2. It felt as if the virtual hand I saw was someone else's
3. It seemed as if I might have more than one hand
4. It felt like I could control the virtual hand as if it was my own hand
5. The movements of the virtual hand were caused by my movements
6. I felt as if the movements of the virtual hand were influencing my own movements
7. I felt as if the virtual hand was moving by itself
8. The virtual hand touched the same part of the disc that my real hand touched
9. The virtual hand's position in the virtual setup corresponded to my real hand's position in the real setup

10. It felt as if my (real) hand was turning into an 'avatar' hand
11. At some point it felt that the virtual hand resembled my own (real) hand, in terms of shape, skin tone or other visual features.

Participants responded in a 7-point Likert scale from -3 (strongly disagree) to 3 (strongly agree).

Ownership score = (Q1 + Q4 + Q9 + Q11)/ 4.

Agency score = (Q6 + Q4)/2

*Ownership and Agency scores correlations with Agency slopes and Mratio:*

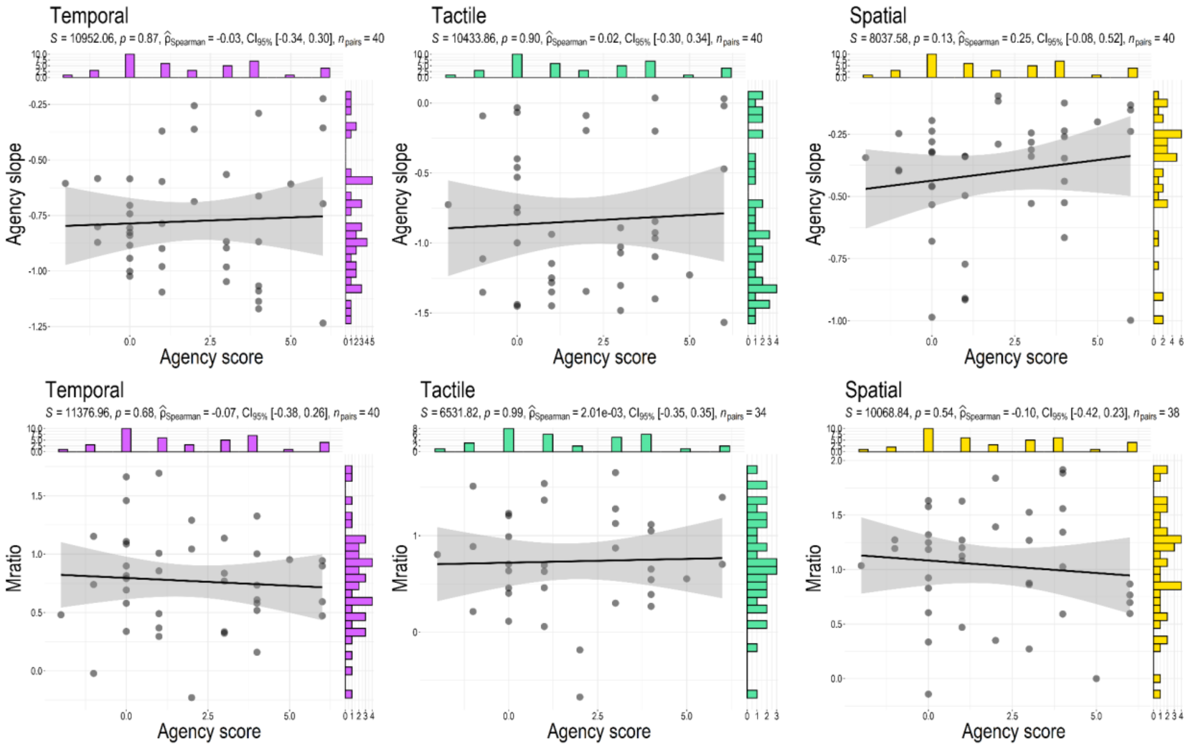


**Fig. S2.** Correlation between the Agency slope and Mratio (metacognitive efficiency) with Agency score as measured with the Embodiment Questionnaire. In all panels, each dot corresponds to one participant, the histograms indicate the distribution of data for each condition, the black lines represent the regression line, and the shaded area corresponds to the 95% confidence interval.

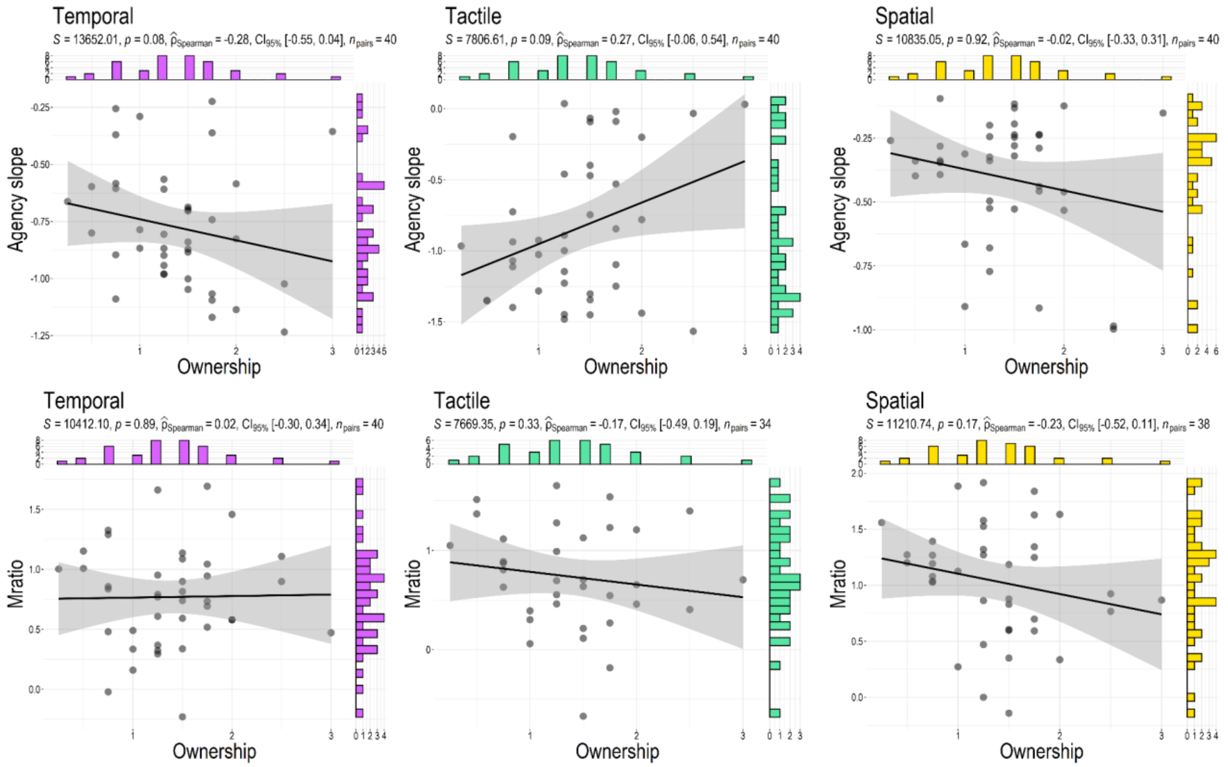


**Fig. S3.** Correlation between the Agency slope and Mratio (metacognitive efficiency) with Ownership score as measured with the Embodiment Questionnaire. In all panels, each dot corresponds to one participant, the histograms indicate the distribution of data for each condition, the black lines represent the regression line, and the shaded area corresponds to the 95% confidence interval.
